# A chance insight into phosgene toxicity

**DOI:** 10.1101/2024.04.23.590734

**Authors:** Ekin Daplan, Enrique Rodriguez, Nick Lane, Luca Turin

## Abstract

It has long been known that phosgene, a war gas and an industrial reagent, causes intense oxidative stress, but how it does so remains unclear. Here we report an accidental discovery: Electron spin resonance spectroscopy (ESR) of live fruit flies reveals that phosgene exposure results in a distinctive manganese (II) hyperfine structure. After exposure to phosgene, every batch of flies consistently displays the Mn (II) signal. Regardless of the aftercare provided, these flies inevitably perish, making the signal a diagnostic of phosgene poisoning in flies. The intensity of the signal is dependent on both exposure time and concentration, resembling the kinetics of phosgene poisoning. The signal of Mn (II) correlates with the presence of a functional superoxide dismutase *Sod2*. After exposure, heterozygous *Sod2* mutants have a markedly lower intensity of Mn (II) in their ESR spectrum. We suggest that phosgene disturbs Mn redox cycling between ESR-silent Mn (III) and ESR-active Mn (II) that is required for superoxide dismutation. Accordingly, mitochondria of phosgene-treated flies show reduced rates of hydrogen peroxide production, and severely compromised complex I-linked respiration. It is likely that phosgene damages mitochondria through MnSOD and complex I, which contributes to its toxicity. This work uses *Drosophila melanogaster* for the first time in phosgene research.

## Introduction

While studying the effect of general anesthetics on the electron spin content of *Drosophila*, we encountered an unexpected phenomenon. Exposing the flies to vapor from an aged batch of chloroform resulted in a striking hyperfine six-line electron spin resonance (ESR) signal, as opposed to the single carbon-centered radical resonance typically observed^1,2^. The flies never recovered from the exposure and exhibited signs of poisoning, among which a phase of spasmodic proboscis movements preceding death. We were unfamiliar with the new ESR signal, but a search for “ESR six lines” revealed it to be diagnostic of manganese in the Mn (II) state^3^. Because chloroform gradually degrades to phosgene over time^4,5^, we suspected that the Mn (II) signal was caused by phosgene.

Manganese is a relatively unusual metal in biochemistry^6^. Several crucial enzymes contain Mn, notably mitochondrial pyruvate carboxylase and superoxide dismutase. MnSOD is a highly conserved enzyme that uses manganese as both an electron acceptor and electron donor in two different reactions, both of which turn the superoxide anion into less reactive species, respectively oxygen and hydrogen peroxide. The manganese cycles between Mn (III) and Mn (II) (Figure 1). With its ability to reduce Mn (III), MnSOD stands out as the sole candidate capable of generating a Mn (II) signal, detectable in X-band ESR. SODs are the main defense against superoxide radicals, potentially harmful reactive oxygen species produced by aerobic respiration^7^. In animals, MnSOD is restricted to mitochondria, where the electron transport chain is located, and the majority of superoxide is produced. The other form of SOD, CuZn SOD, is present in the cytosol only. The quantity of MnSOD in tissues varies depending on the abundance of mitochondria^8^. Interestingly, oxygen toxicity, presenting similar symptoms to phosgene poisoning^9^, is more severe in MnSOD deficient mutants ^10^. MnSOD expression is tightly regulated. While tissue-specific overexpression might introduce some benefits^11–13^, both extremes of expression levels are reported to cause severe problems in mice and fruit flies^14,15^.

**Figure 1.**
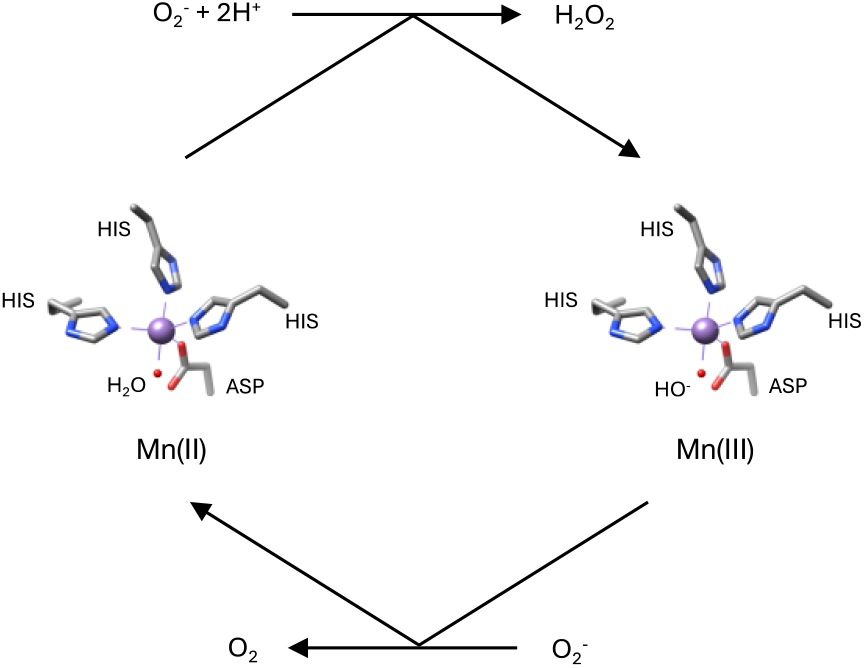
The superoxide dismutase activity cycle, redrawn from^16^

Phosgene, also known as carbonyl dichloride, was discovered and named by John Davy in the early 19^th^ century^17^. In World War I it was a notorious warfare agent, believed to exert its poisonous properties by producing HCl in the presence of water to irritate the respiratory track^18^. Fortunately, it is no longer employed as a war gas, but it continues to serve various industrial purposes. It is now known that due to its low water solubility, phosgene does not induce irreversible irritation reacting with the mucous surfaces in the respiratory tract. It travels to the lower areas of the lungs and possibly passes through the air-blood barrier^19,20^. The effects of exposure to high concentrations of phosgene typically follow three phases. The first is an immediate, transient choking reaction, likely due to the release of hydrochloric acid on contact with water and lung tissue. The second is a symptomless delay phase, of length inversely proportional to the exposure dose of gas, and typically lasting hours^21^. The third phase begins when the deleterious effects of phosgene begin to manifest themselves. Fluid leaks into the lungs, breathing becomes impaired, and death can follow rapidly. Phosgene poisoning is uncommon, the mechanism of action is unknown, there is no cure, and it is not possible to predict or prevent the onset of its distinct physiological failures^22–24^.

The only stable isotope of manganese possesses nuclear spin, enabling certain redox states to exhibit a hyperfine structure in ESR. Here we demonstrate that exposure to phosgene triggers a transient Mn (II) signal in the ESR spectrum of live flies. The signal is detectable for approximately 17 hours following the exposure. The intensity of the Mn (II) signal increases with either increasing duration of exposure or with the concentration of phosgene. These kinetics, referred to as Haber’s rule^25^, are considered to be an indicator of the severity of intoxication by several poisonous gases, including phosgene^26^.

Phosgene poisoning involves the failure of various biological processes^23^. The physiological complications involving free radicals and their clearance are difficult to ascribe to a single molecular dysfunction. In this article, we aim to establish whether MnSOD redox cycling is disturbed by phosgene and is an important player in phosgene poisoning. We show that the presence of phosgene-induced Mn (II) spin signal in fruit flies depends on *Sod2* expression. We performed molecular dynamics to calculate the likelihood of phosgene reacting with tyr34, a site proven to be important for MnSOD functioning^27^. MnSOD is localized in mitochondria, responsible for converting the relatively harmful superoxide radical to less harmful hydrogen peroxide. The absence of a functional MnSOD in mutant mice is reported to produce catastrophic complications involving the inhibition of respiratory complexes^28^. Therefore, in our final step, we analysed mitochondrial function following phosgene exposure. Using high-resolution fluorespirometry, we measured oxygen flux and hydrogen peroxide concentration in a range of specific respiratory states. Our findings show that flies subjected to phosgene do indeed generate less hydrogen peroxide in conditions where superoxide flux from the respiratory chain is maximized. Moreover, the activity of complex I was significantly impaired after phosgene poisoning.

## Methods

### Drosophila melanogaster

Flies were cultured using an instant *Drosophila* mix obtained from Advanced Husbandry. This mixture was supplemented with propionic acid and nipagin in order to postpone spoilage and protect the medium from unwanted organisms. The ambient temperature was 18-25 °C with a humidity range of 40-80 %. The natural light cycle of the Midlands United Kingdom was followed. Our wild-type flies were the Oregon-R strain, obtained from Darwin Biological. We used y^1^w^1^ for control (Bloomington *Drosophila* Stock Center #1495), and two heterozygote mutants with deletion of *Sod2* coding region, *Sod2*^*Δ2*^ and *Sod2*^*Δ12*^ (Bloomington Drosophila Stock Center # 27643, # 27644)

### Phosgene Aliquots Preparation

Phosgene solution was purchased from Sigma-Aldrich. The aliquots of 25 μl, 50 μl, and 100 μl were pipetted into a 1.1 ml glass screw-top autosampler vial. In addition to their own lid, the vials were supplemented with a Teflon lid for the protection of the plastic from phosgene. All aliquots were kept at +4 °C to avoid evaporation of phosgene.

### Modified Micro Test Tube

The test tube (A, Figure 2) was selected due to its compatible dimensions with a 1.1 ml screw-top tube (B, Figure 2) and its durability against phosgene. The tube bottom was cut off and plugged with a shortened pipette tip wrapped with a 225-micron nylon mesh purchased from Plastok. A snug fit was obtained, and excess mesh trimmed. The flies then sit a few mm above the tube bottom.

**Figure 2.**
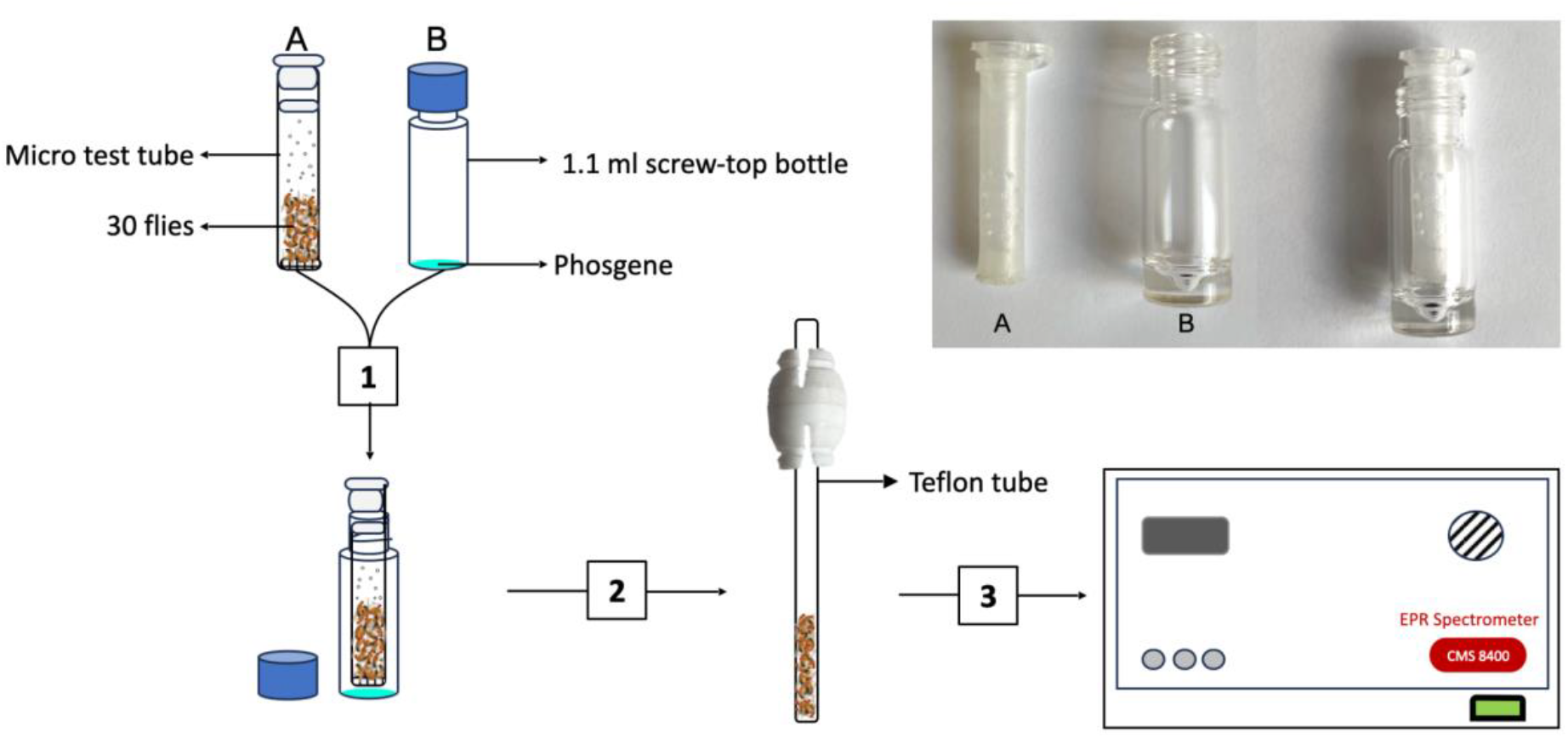
Experimental steps are depicted in the figure.

### Phosgene Exposure

The 1.1 ml glass screw top autosampler vial that contains phosgene was freshly taken from +4 °C and placed under the fume cupboard at room temperature. The vial was opened immediately, and flies in a Teflon tubing were placed inside the micro test tube, then the test tube was placed inside the bottle. They remained for the required duration of exposure (1, Figure 2). After exposure, the flies were tapped inside the teflon tube (2, Figure 2). Lastly, the teflon tube with flies was rested inside the spectrometer for the measurement (3, Figure 2).

### ESR Protocol

ESR spectroscopy on live flies has been described before^1,2^. In our protocol, the control ESR trace was obtained from 30 male fruit flies inside a PTFE tube. The spectrum was measured before and after the exposure.

CW (continuous wave) ESR was used for the spin measurements. All the parameters were identical for control and mutant strains. The central field was set to 3370 gauss with an 800 gauss width. Ambient gas inside ESR was a constant flow of nitrogen gas to keep flies immobile. Adult *Drosophila melanogaster* are known to survive anoxia for up to 12 hours^29^. The experiments were conducted at room temperature.

### Mitochondrial function analysis through high-resolution fluorespirometry

Simultaneous measurements of oxygen consumption and H_2_O_2_ flux in various respiratory states were performed on permeabilized thoraces, and on isolated mitochondria from whole male flies at 23 °C using an O2k-FluoRespirometer (Oroboros Instruments, Innsbruck, Austria) as described previously^30,31^. Three permeabilised thoraces of control or phosgene-exposed flies were transferred to a multi-welled plate containing 2 ml of ice-cold preservation solution BIOPS (2.77 mM CaK2EGTA, 7.23 mM K2EGTA, 6.56 mM MgCl2·6H2O, 20 mM imidazole, 20 mM taurine, 15mM Na2 phosphocreatine, 0.5 mM dithiothreitol, 50 mM K-MES, 5.77 mM Na2ATP) and 81.25 µg/ml saponin for permeabilization for 20 minutes, after which they were rinsed for 5 min in 2 ml of MiR05 respirometry buffer (0.5 mM EGTA, 3 mM MgCl2.6H2O, 60 mM lactobionic Acid, 20 mM taurine, 10 mM KH2PO4, 20 mM HEPES, 110 mM D-sucrose, 1 g/L BSA, pH 7.1) before their wet weight was recorded, and transferred to the O2k chamber. Isolated mitochondria from 20 whole flies were used following previously published methods^32^.

Oxygen and fluorescence signals were calibrated daily as per the manufacturer’s protocols. For H_2_O_2_ analysis, 15 µM DTPA, 1 unit HRP and 10 µM AUR were injected sequentially in the chamber prior to sample addition. After the permeabilised thoraces or isolated mitochondria were added to the chambers, and a substrate-uncoupler-inhibitor titration (SUIT) protocol was conducted as follows: first, NADH-pathway substrates pyruvate (10 mM) and malate (2 mM) were added, and the signal was left to stabilise for 15 minutes to obtain N-LEAK state respiration (LEAK). Then, ADP (5 mM) was added to reach the N-OXPHOS state (CI-OXPHOS). Sequential additions of proline (10 mM, CI+Pro), succinate (10 mM, CI+Pro+CII), glycerophosphate (10 mM, CI+Pro+CII+Gp, Max respiration) then followed, and maximal coupled respiration rates were obtained. Titration with the uncoupler FCCP in 0.5 µM increments allowed the estimation of the maximum uncoupled respiration (Max uncoupled). The N-pathway was then inhibited of with rotenone (0.5 µM, Rotenone-inhibited), the S-pathway with malonate (5 mM, Mn-inhibited), and finally complex III with antimycin A (2.5 µM, Antimycin A-inhibited), allowing the estimation of residual oxygen consumption (ROX). Chambers were opened for reoxygenation and closed before injection of ascorbate and TMPD (2.5 mM and 0.5 mM respectively) to measure complex IV oxygen consumption, after which the enzyme was inhibited by the injection of 100 mM of sodium azide to calculate complex IV activity (CIV, corrected for chemical background as per the manufacturer’s instructions). In a series of separate experiments, we measured H_2_O_2_ flux in reverse electron transfer conditions (RET), by the successive additions of glycerophosphate (10 mM, glycerophosphate-RET), and antimycin A (2.5 µM, Antimycin A-RET). A final addition of CuZn SOD (5 U/ml) was performed to evaluate the amount of undetected H_2_O_2_ released.

### Statistical Analysis

ESR experiments with mutants were carried out three times with three distinct fly populations. The significance was determined using the intensity difference in the Mn (II) hyperfine structure. The magnetic field at each peak and trough was noted, and the average of eight experiments was used to standardize the location and corresponding value per signal (Figure SM1). The difference between peak and trough was referred as intensity. We used the intensity of 5 peaks out of 6 in the Mn (II) signal to quantize the difference between the control and experimental groups. The significance was determined using Welch’s t-test in R.

O_2_ and H_2_O_2_ fluxes were extracted using the DatLab 7.4 software, processed using the manufacturer’s calculation templates, and analysed by correcting by either thorax wet weight, or protein content of the isolated mitochondria (using the QuantiPro BCA Assay Kit, Sigma-Aldrich). We calculated respiration-associated parameters as follows: OXPHOS coupling efficiency = (LEAK/CI-OXPHOS)*100; E-T coupling efficiency = LEAK/Max uncoupled; substrate contributions as the fractional change in flux upon addition of the substrate (proline, succinate glycerophosphate contr.); uncoupling effect = (Max uncoupled/Max respiration) *100; Excess ETS capacity = 1-(Max respiration /Max uncoupled); Complex I contribution = (Max uncoupled – Rotenone-inhibited)/ Max uncoupled *100; Complex I control ratio = (CI-OXPHOS / Max respiration) *100. Boxplots and statistical analyses were done in R using ggplot2. The differences in oxygen or H_2_O_2_ flux at each state or parameter were evaluated through a two-sample Welch’s t-test in R.

## Results

### Phosgene gives rise to Mn (II) signal in the ESR spectrum of fruit flies

Figure 3 depicts our main finding. The left trace shows the ESR spectrum of 30 live flies (approx. 30 mg, see methods). The central resonance is caused by stable free radicals found in fruit flies, specifically melanin^33^. The right trace depicts the same flies following a 10-minute exposure to phosgene diluted in toluene (see methods). Two striking changes developed in the ESR spectrum after phosgene. The central melanin resonance is significantly reduced (1), while six prominent hyperfine lines have appeared that overlap with it (2). The six-line pattern identifies Mn (II) ions in solution^34^. A control with only toluene shows no hyperfine lines (Figure SM2).

**Figure 3.**
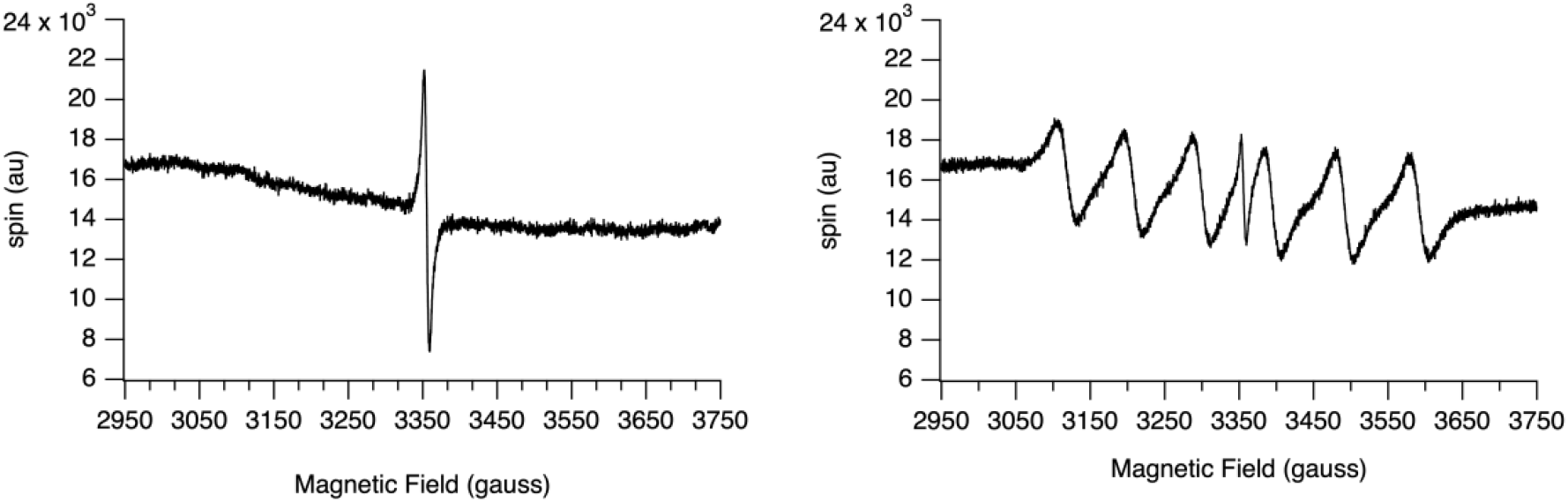
Two X-band scans of 30 fruit flies ten minutes apart. The left scan is the control, showing a typical carbon-centered radical signal at 3356 G, approximately 10 G in width. The right scan shows the same flies after a 10-minute exposure to phosgene vapor. The carbon-radical signal is reduced in amplitude, and six new equally spaced lines have appeared, diagnostic of Mn (II).

### The Mn (II) signal requires metabolism

After phosgene exposure, flies cannot recover, but exhibit small, repetitive movements, indicating that they are still alive (Figure SM4). To begin, we wondered if the signal we observed was simply due to a chemical process. To test this, we exposed dead flies to phosgene. Unlike living flies, dead flies did not show the six characteristic peaks, but they did show a small drop in the central melanin signal (Figure SM3). We deduce that whatever was needed to produce Mn (II) was not present or working properly. Fly cuticle is rich in melanin. Phosgene’s first encounter with flies is through their cuticle. The majority of the radical content of melanins and sclerotins comes from their semiquinone and quinone radicals, which are both derived from phenols. Phosgene reacts with phenols to produce chloroformates, which are then unable to form quinones. The slight reduction of the melanin resonance, therefore, can be accounted for by chloroacylation of phenols by phosgene.

### The Mn (II) signal is transient

Remarkably, the effect of phosgene on both features of the ESR spectrum is temporary. The ESR spectra for Mn (II) and melanin content remain unaltered for ∼1 hour, after which the Mn (II) signal gradually disappears while the melanin signal gradually returns (Figure 4).

**Figure 4.**
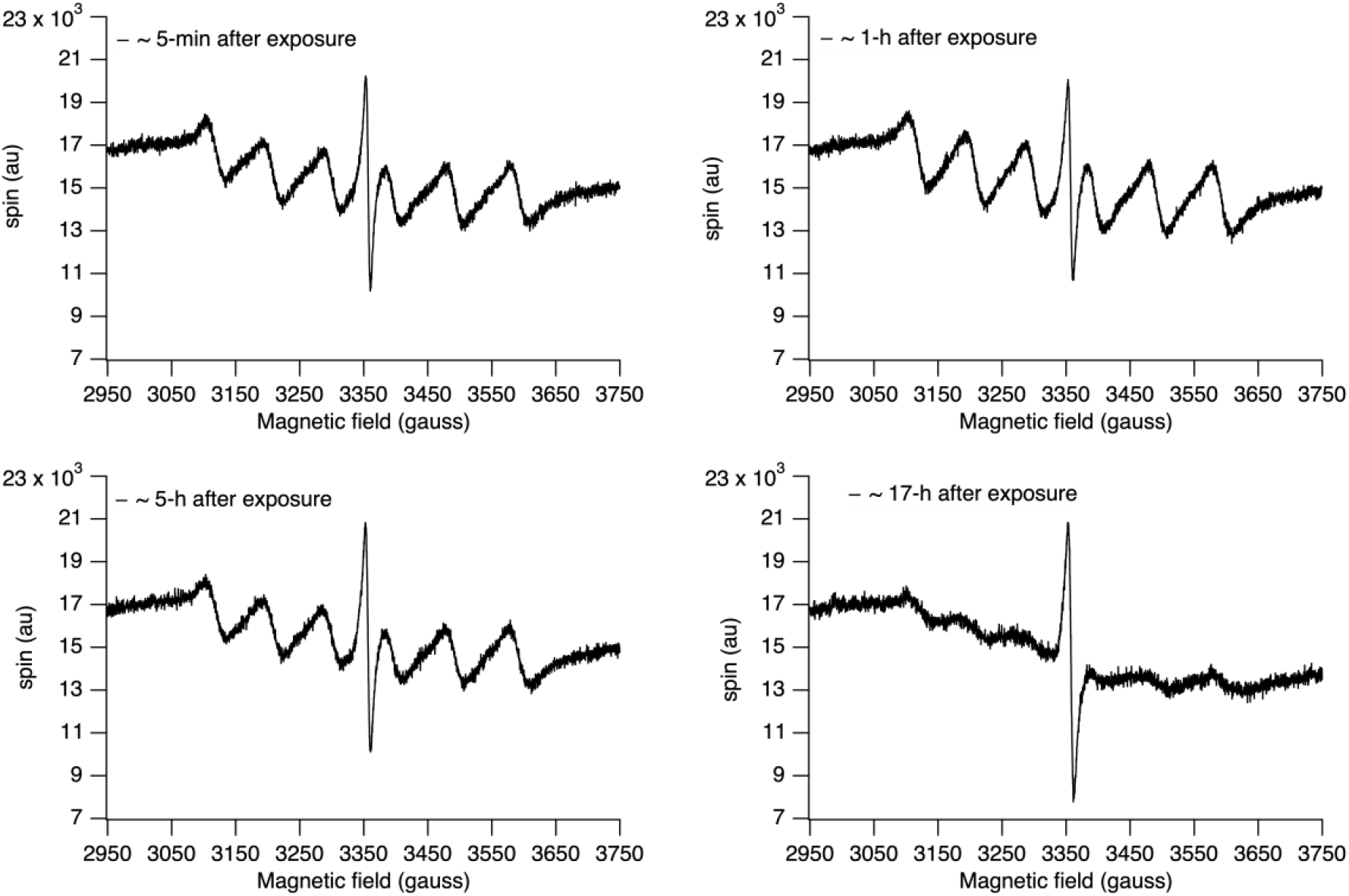
The Mn (II) and melanin peaks as a function of time after a brief phosgene exposure. ESR traces taken at different times after a single 10-minute exposure to phosgene vapor. The Mn (II) signal appears rapidly, then decreases slowly over hours.

### Mn (II) signal intensity depends on both the duration and dose of the phosgene exposure

While experimenting with various doses of phosgene, we discovered that there was a limit to the intensity of the signal, and increasing the dosage did not boost the signal further. On the other hand, a small amount of phosgene was enough to cause the same intensity of the signal when applied for a prolonged period of time. In order to examine this phenomenon closer, we exposed flies to 25 μl, 50 μl, or 100 μl of phosgene for 5-min, 10-min, or 20-min (Figure 5).

**Figure 5.**
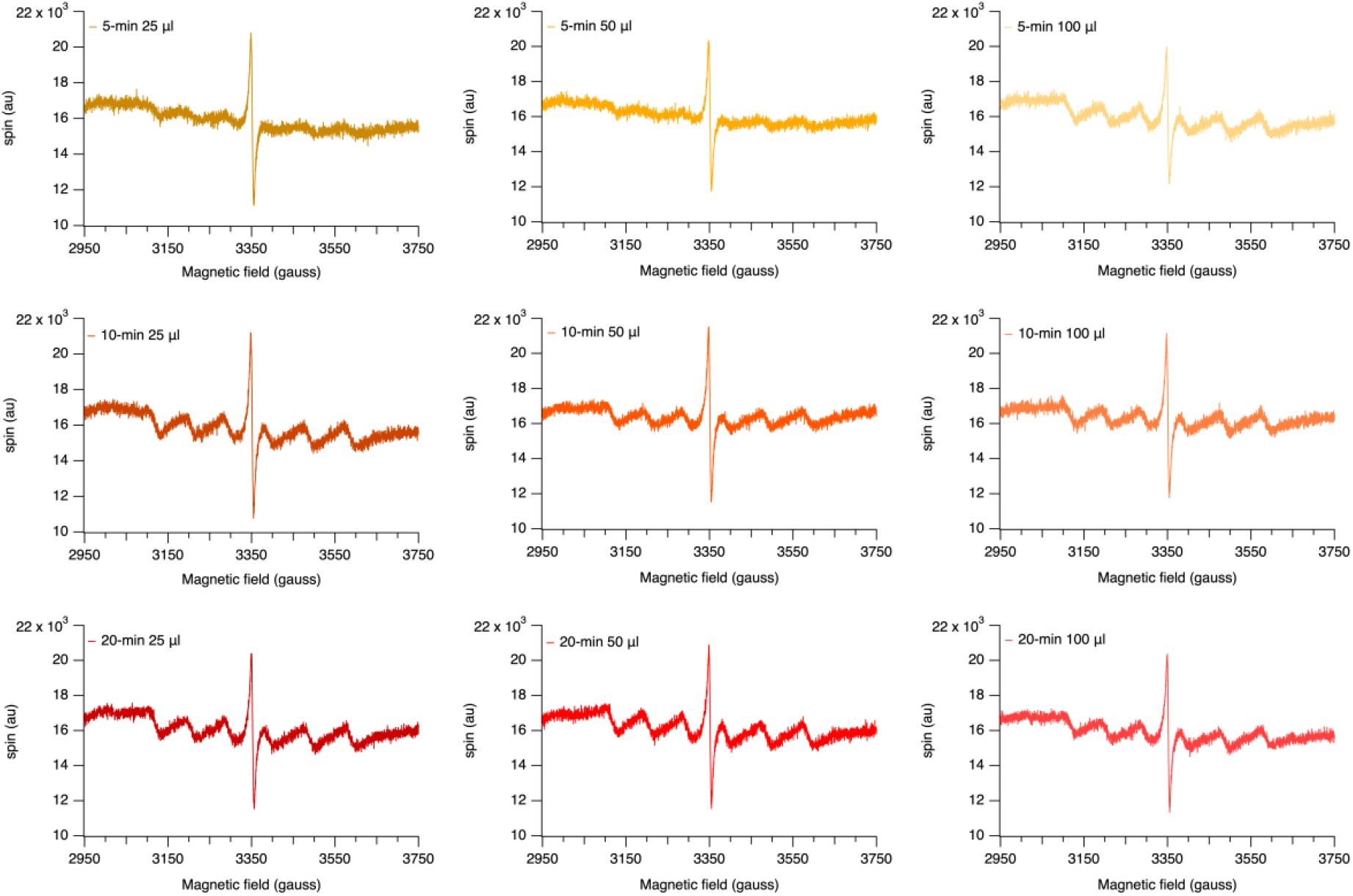
ESR spectrum after phosgene exposure is shown in the figure by increasing in dose from left to right and increasing time from top to bottom.

### Signal intensity is lower in MnSOD mutants after phosgene exposure

A direct test of the involvement of MnSOD in the Mn (II) signal is provided by mutants with only one functional *Sod2* gene (Figure 6). Two of these (Sod2^Δ02^ and Sod2^Δ12^) are available, in which the coding region of one allele of the *Sod2* gene has been deleted with P-element activity leading to complete loss of function. The mutants are in a yellow-white (yw^1^) background, and therefore controls must be performed in the same background for comparison. For the experiment, the protocol was followed; Sod2^Δ02^, Sod2^Δ12^ and yw^1^ flies were subjected to 25 μl of phosgene for 10 minutes. The difference in the ESR spectrum of the control and experimental groups was noticeable with the naked eye. We quantified the difference by measuring the intensity of each Mn (II) peak per experiment. Both mutants showed significant differences compared with the controls (p<0.001). The sample size for the three groups was unequal because of the challenges in cultivating the mutant flies, particularly Sod2^Δ12^. The experimental variation was quite high in the control group.

**Figure 6.**
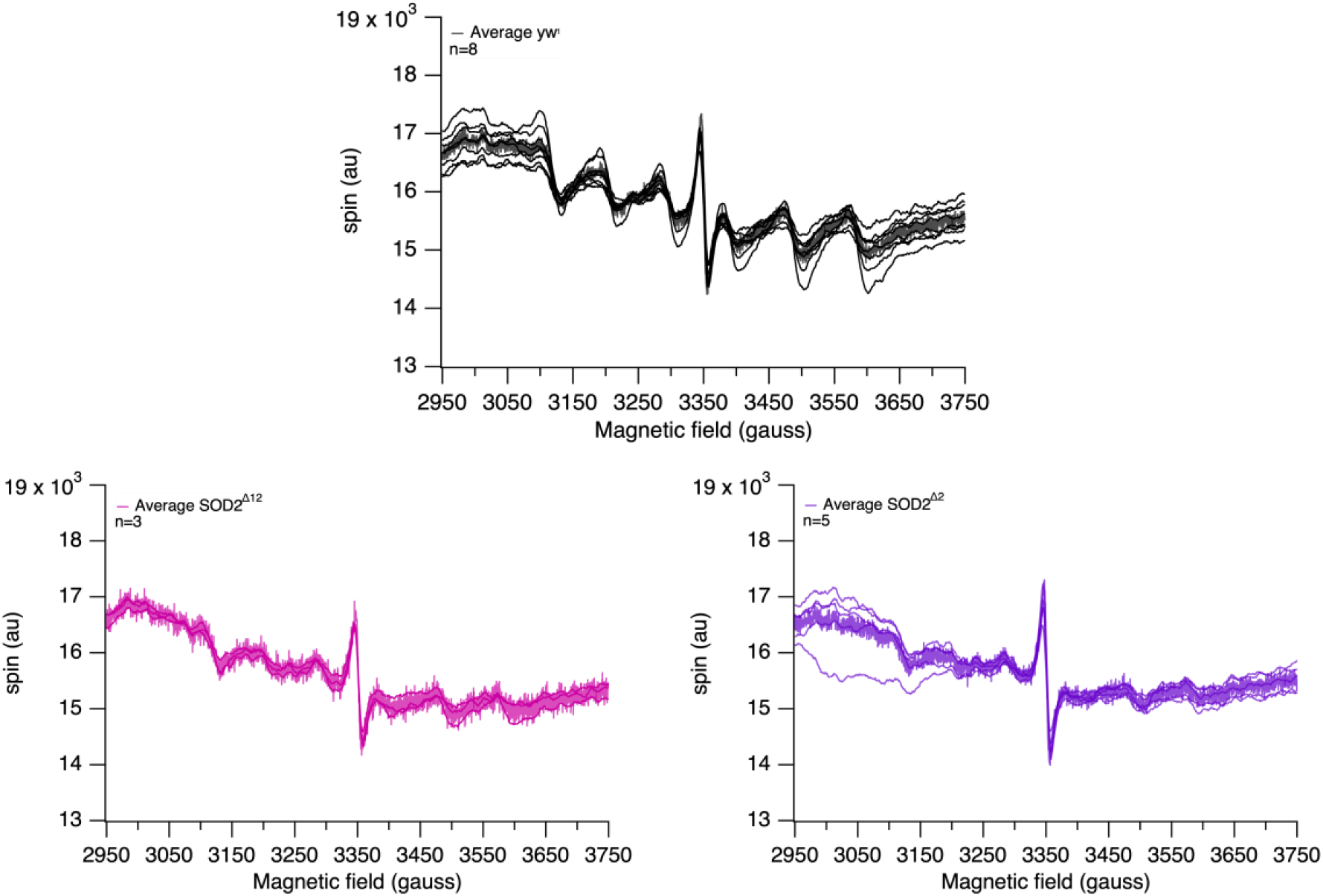
ESR spectrum of experimental and control groups are plotted. SOD2^Δ12^ is represented by pink colour, and SOD2^Δ02^ purple. The p-value for both mutants is <0.001 when compared to the control.

### The signal intensity depends on the fly’s age

Given that the experiments controlled for variations in time and dose of exposure, we questioned the potential source of the remaining variation. We replicated the experiments with age-matched fly groups for yw^1^ and Sod2^Δ2^. We chose not to use Sod2^Δ12^ again, given our challenges in cultivating the flies; we also wanted to minimize the usage of phosgene in the laboratory. The results revealed practically no detectable Mn (II) at 1-day Sod2^Δ2^, and a definite, though smaller Mn (II) signal at day 7 (Figure 7). The difference in peak amplitudes between control and *Sod2* mutant is statistically significant on both 1-day and 7-day flies. However, according to Flybase data, the expression level of *Sod2* does not vary with developmental stage, with expression levels of *Sod2* being high throughout the life of a fly^35^.

**Figure 7.**
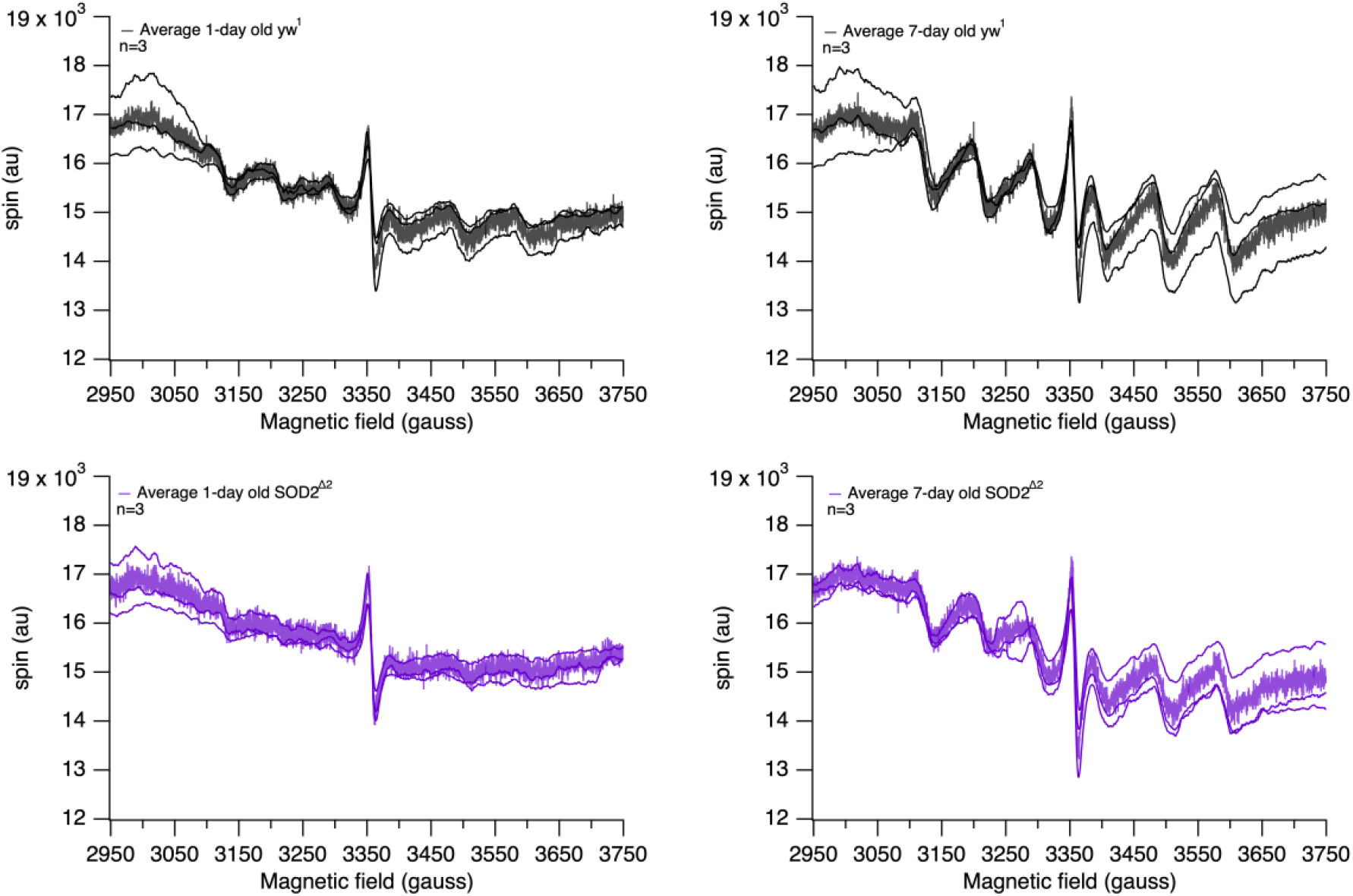
ESR spectrum of experimental and control groups are plotted. Black lines represent the yw^1^ and purple SOD2^Δ2^ mutant. On the left, spectrum of 1-day old flies and on the right 7-day old flies are shown. P-value was calculated by measuring intensity of each peak and comparing control and experimental for both 1-day and 7-day old flies. P-value between each graph above has a p-value <0.001.

### Chloroformylation of MnSOD by phosgene is feasible

Carbonyl dichloride is an acylating agent that reacts with hydroxyl groups and other nucleophiles to form chloroformates^4^. The reaction with phenols is of primary interest here, because phenols are both abundant in melanins and present in proteins as tyrosine side-chains. Furthermore, a highly conserved tyrosine (tyr34) is located near the SOD2 active site and is necessary for SOD2 function^27^ (Figure 8; Figure SM5). We therefore hypothesize that the Mn(II) signal is caused by chloroacylation of Tyr 34 in SOD2 (equation 3), which prevents the Mn ion from cycling, a key part of its catalytic mechanism (equations 1 and 2)^36^. We cannot, of course, rule out the possibility that acylation occurs on another amino acid, for example nearby serine 131, also conserved, and that the effect of acylation on Mn redox cycling is caused by a change in conformation. The recovery may arise from hydrolysis of the ester produced in reaction 3 to regenerate the phenol^37^. It is less likely to be caused by reaction (4) above, which would require 4-chlorotyrosine to allow SOD2 to function, and chlorophenols to form quinone radicals.

**Figure 8.**
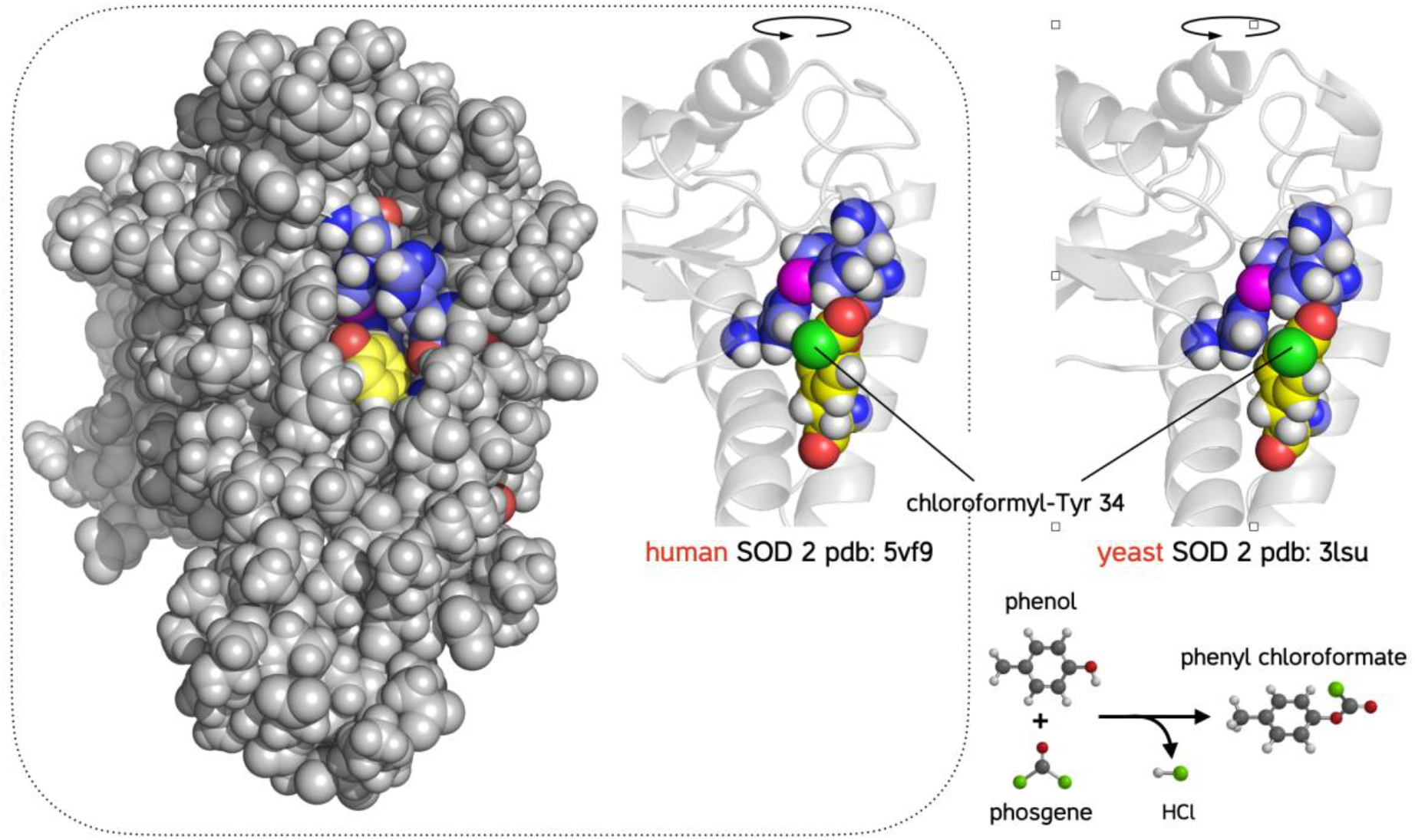
Left: All-atoms model of human mitochondrial superoxide dismutase 2. The pdb model (5vf9) was imported into a molecular mechanics code (see Methods), hydrogens and solvent water (not shown) were added and the structure energy-minimised. The tyrosine 134 ring is shown in yellow, with phenolic oxygen visible in red, the histidines surrounding the manganese ion in blue and the manganese ion itself in pink. Inset, middle: The tyrosine was chloroformylated and the resulting structure energy-minimised again. The chloroformyl group fits in the active site. Inset, top right: The active site in *Saccharomyces cerevisiae* SOD 2, pdb 3lsu after energy minimisation. The structure of the two enzymes are very similar.

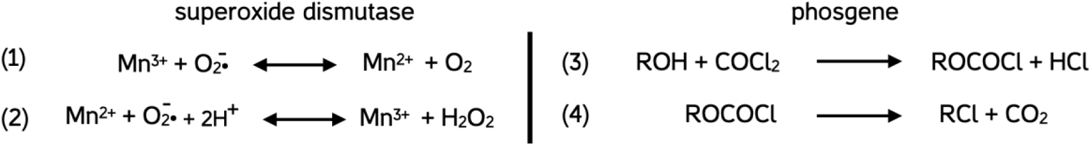

To evaluate the likelihood of reaction of phosgene with SOD, we performed a molecular mechanics energy minimisation on the all-atoms human MnSOD structure (pdb 5vf9) in water (Figure 8). The tyrosine (yellow) is exposed to the solvent, right next to the three histidines (in blue) coordinating the Mn ion. Chloroformylation of the tyrosine followed by energy minimisation causes only a small outward movement of the tyrosine, shown at right in human and bacterial MnSOD, whose structures are highly conserved. It therefore seems plausible that chloroformylation of MnSOD can occur, and likely disrupt MnSOD function. The importance of try34 on the function of MnSOD has been reported before^27^.

### Mitochondrial respiration and H_2_O_2_ efflux are severely compromised after phosgene exposure

To assess the impact of phosgene on mitochondrial functions, and particularly on H_2_O_2_ flux, we used high-resolution fluorespirometry to compare responses in the thorax and in isolated mitochondria of control and phosgene-exposed OreR *Drosophila* (Figure 9).

**Figure 9.**
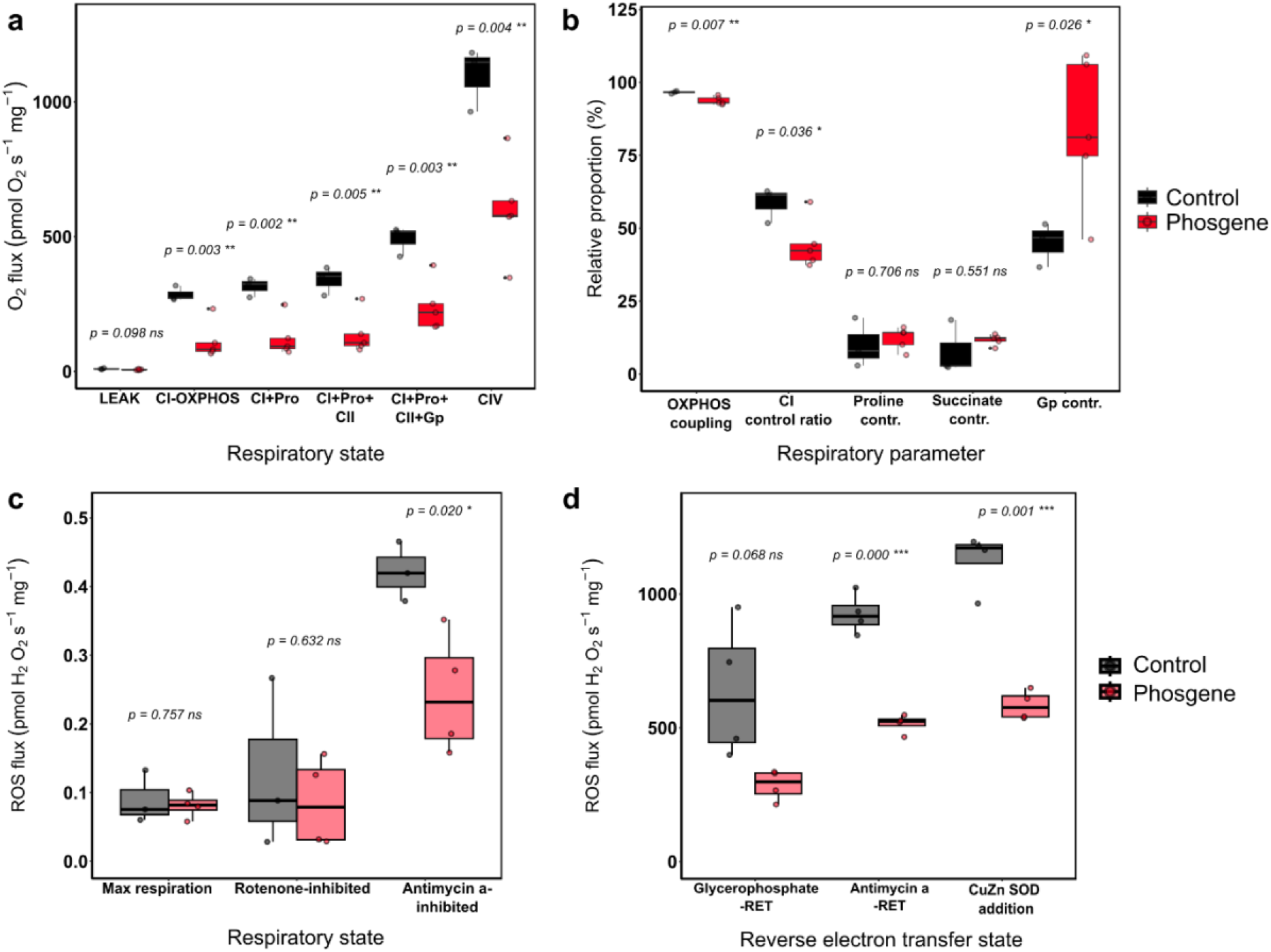
Mitochondrial respiration (a, b) and reactive oxygen species flux (c, d) in control and phosgene-exposed flies. (a) O_2_ flux measured in successive steps of a SUIT-protocol in permeabilized thoraces, and (b) detailed functional parameters calculated from these measurements. See methods for detailed explanations of all states. (c) ROS flux as hydrogen peroxide detected in permeabilized fibers, in the same SUIT-protocol; and (d) in isolated mitochondria in a protocol analysing the reverse electron transfer condition.

Respiration coupled to phosphorylation (OXPHOS) with complex I substrates was suppressed by more than 50% (*p* = 0.003), as was maximal respiratory flux with a combination of substrates targeting complexes I, II, III and proline and glycerophosphate dehydrogenases (*p* = 0.003). The maximal capacity of the terminal electron acceptor complex IV was also decreased to about 45% of that in control flies (*p* = 0.004, Figure 9a). We then dissected the impact of phosgene exposure on the different respiratory pathways (Figure 9b). We found significantly reduced OXPHOS coupling efficiency, i.e. the coupling of electron transfer to phosphorylation upon addition of ADP (*p* = 0.007), indicating compromised respiration with complex I substrates. This was further confirmed by the 25% decrease in complex I control ratio (*p* = 0.036), meaning that the relative contribution of this complex to maximal O_2_ flux was severely affected in flies exposed to phosgene, while the contribution of substrates proline (ProDH) and succinate (complex II) was not significantly different between the two groups (*p* = 0.706 and *p* = 0.551, respectively). On the other hand, the contribution of glycerophosphate to respiration, was around 45% higher in phosgene-exposed flies compared to controls (*p* = 0.026), suggesting that the mG3PDH and complex III were less affected. .

We simultaneously measured the flux of reactive oxygen species, corresponding to the efflux of H_2_O_2_ from the permeabilized tissues (Figure 9c), and found no differences during maximal coupled respiration (*p* = 0.757) or when inhibiting respiration at complex I with rotenone (*p* = 0.632). However, when stimulating maximal ROS production with antimycin a, phosgene-treated flies had a 40% reduced H_2_O_2_ flux (*p* = 0.02). We then probed the maximal capacity of ROS efflux by stimulating reverse electron transfer in isolated mitochondria (Figure 9d) from whole flies and found H_2_O_2_ flux to be around 45% reduced with glycerophosphate alone (*p* = 0.068), and 55% reduced when adding antimycin-a (*p* < 0.001). Subsequent addition of CuZn SOD to the system further increased ROS flux by approximately 15%, but the difference between the two groups remained consistent at around 50% (*p* = 0.001).

## Discussion

In this study, we documented potentially important biochemical aspects of phosgene poisoning, which we discovered after detecting a Mn (II) signal in the ESR spectrum of fruit flies upon exposure to phosgene (Figure 3). We reasoned that the appearance of Mn (II) in the spectrum must be a sign of malfunctioning MnSOD because there is no possibility of Mn itself being introduced by phosgene exposure. The Mn (II) signal must arise from the reduction of ESR-silent Mn (III) present in the flies, somehow caused by phosgene. We further speculated that defective MnSOD might lead to poisoning the animals. There are several reasons for us to suspect this. Firstly, none of the flies survived after the displayed Mn (II) signal, emphasizing the correlation between Mn (II) and phosgene lethality. Secondly, MnSOD is the most critical cellular defence mechanism against reduced oxygen species among SODs. Organisms cannot afford deletion of *Sod2*. Halliwell notes that “transgenic animals lacking MnSOD are the most dramatic cases among SODs”^8^. Inactive MnSOD therefore seemed a plausible explanation of the severity of phosgene poisoning. We tested this idea in two different ways. First, we changed the phosgene dose and the amount of MnSOD in the flies to see whether we can alter the intensity of the Mn (II) signal. We demonstrated that increasing the concentration or duration of phosgene exposure could result in an increase in Mn (II) intensity in the ESR spectrum of flies, mimicking the kinetics of phosgene poisoning (Figure 5). Furthermore, we investigated the differences in Mn (II) signal intensity between mutant lines with only one functional *sod2* gene. We discovered that the mutants had reduced signal intensity, indicating that MnSOD is involved in the production of Mn (II) signal in flies (Figures 6 and 7). Second, we used fluorespirometry to investigate phosgene’s effects on mitochondria. Because MnSOD is responsible for converting superoxide to hydrogen peroxide, we measured the hydrogen peroxide levels of flies after phosgene exposure. We found less hydrogen peroxide production (Figure 9), which is consistent with the problem being a lack of functional MnSOD. Furthermore, phosgene-treated flies showed decreased activity in their mitochondrial complexes I and IV (Figure 9). The impairment of respiratory complex I and consequently depressed respiration rate could be due to a direct effect of phosgene on this enzyme, or to increased superoxide levels not subsequently detoxified. The interactions between the activity of complexes, and different reactive oxygen species are likely to involve MnSOD but need further work to be untangled^38^. The correlation between functional MnSOD and reduced mitochondrial complex activity has previously been reported^28^.

Currently, oxygen treatment is considered a main intervention in the treatment of phosgene poisoning, particularly in symptomatic patients^39,40^. However, research into this issue paints a more complex picture. For example, in a study to examine the role of oxygen therapy in treating phosgene poisoning, it was shown that dogs treated with 95%, 90%, and 40% oxygen after phosgene exposure had a 5-20 % higher short-term survival rate compared with dogs that did not receive oxygen treatment^41^. Curiously, this effect only lasted for the first 3 days in which the oxygen delivery was provided. When the oxygen delivery was stopped, the survival rate of the experimental group sharply decreased and surpassed that of the control group. The experimental group receiving 95% oxygen showed the most significant difference from the control group. While around 9 out of 25 untreated individuals survived, only 2 out of 25 individuals survived in the oxygen treatment group.. Our finding on the decreased activity of complex I and MnSOD function after phosgene poisoning does not provide a definite explanation for the impact of oxygen therapy, but it is possible that hyperoxia might enhance oxygen flux through damaged complex I, temporarily boosting survival, while having no effect on phosgene poisoning of MnSOD. On the contrary, hyperoxia may have exacerbated oxidative stress caused by inhibition of MnSOD. It is important to mention that MnSOD knockout mice were shown to be more susceptible to hyperoxia, and mice overexpressing MnSOD in their lungs were found to exhibit increased resistance to 90% oxygen^10,12^.

Phosgene is an exemplar of Haber’s rule, which relates the dose of a noxious chemical to its effects^25^. In Haber’s analysis, dose means (concentration) x (duration). Remarkably, Haber’s rule applies even to very long exposures at very low concentrations. This raises the possibility that even the low exposure levels in industrial uses of phosgene, commonly used in production chemistry to replace hydroxyls with chlorine^42^, may nevertheless result in lung damage over time^23,43^. The adherence of phosgene to Haber’s rule, and the nature of phosgene as an acylation reagent has long been taken as an indication that the toxicity of phosgene is attributable to the acylation of one or more protein components of lung tissue. If (a) the reaction is irreversible, (b) the damage done is proportional to the fraction acylated, and (c) the concentration of phosgene is the rate-limiting step in the acylation reaction, then Haber’s rule follows. Phosgene acylates nucleophilic amino acid side-chains, including hydroxyls (ser, thr, and tyr), amines (lys) and thiols (cys). Most proteins contain all these amino acids, so this does not usefully narrow down the range of possible targets for phosgene action. Accordingly the action of phosgene has been variously attributed to its effects on glutathione levels, lipid peroxidation and cellular ATP levels^40,44–46^. There is, however, an aspect of phosgene action which may help narrow the range of possible targets. While acylation will proceed in all parts of the body exposed to phosgene, the damage done is concentrated in the lungs. They are, of course, exposed to the highest concentration of inhaled phosgene but also to the highest concentration of oxygen of any tissue of the body. The relation between free radical production, and in particular reduced oxygen species, and phosgene toxicity has been noted many times^47^. Phosgene exposure depletes cellular pathways involved in the cellular response to free radicals^48^. Further, the toxic effects of the inhalation of pure oxygen, which also follow Haber’s rule, are similar to those of phosgene inhalation^49,50^, pointing directly to a role for reduced oxygen species in both cases. Our accidental discovery of the effect of phosgene on MnSOD suggestively brings these threads together: MnSOD is the primary defense mechanism against ROS created during oxidative phosphorylation^51,52^. MnSOD contains a conserved tyrosine hydroxyl close to the active site^53^. That tyrosine has been shown to be crucial to SOD function^27^.

Given the likely large number of cellular components potentially acylated by phosgene, it is unlikely that MnSOD is the only target; therefore, more direct biochemical studies are needed, for example to assess whether the effect on complex I is direct or Tyr34 is unusually reactive. Treatment of tissues with phosgene could yield a rapid and quantitative test for MnSOD. If confirmed by further studies, the mechanism we propose may eventually lead to a rapid diagnostic test of phosgene poisoning, provided the Mn (II) ESR signal due to MnSOD acylation can be detected in other tissues of the body, e.g. blood or skin. We suggest *Drosophila* could be a useful animal model for the development of both tests and therapeutics of phosgene toxicology.

Author’s contributions: ED and LT devised the project, ED performed the ESR experiments, ER performed the O2k Oroboros experiments. ED, ER and LT analyzed data, ED and LT wrote paper, ED, ER, NL, and LT revised paper.

## Acknowledgments

This work was supported in part by a grant from Ionis Pharmaceuticals. Dave Ecker is thanked for his support. ED would like to thank Efthimios Skoulakis for the guidance on the use of *Sod2* mutants.

## Supplementary Materials

**Figure SM1:**
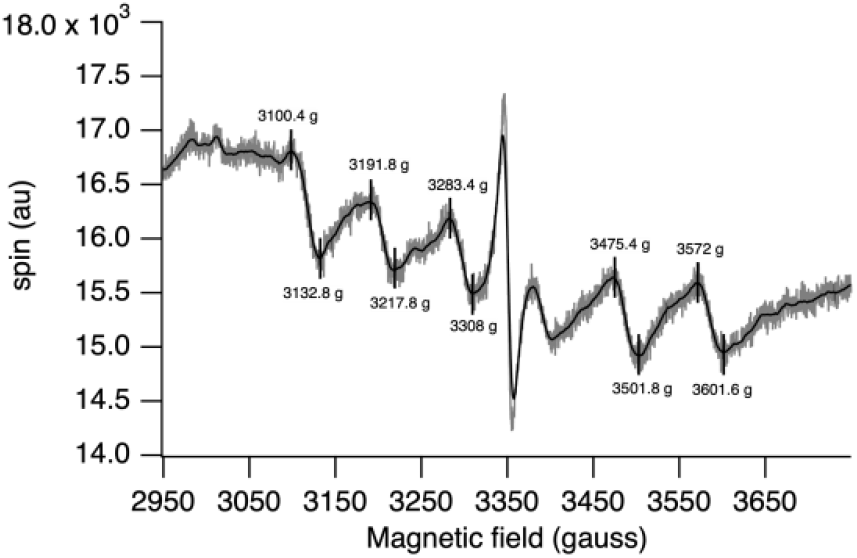
Location of each peak and trough is calculated for the average of 8 experiments.

**Figure SM2:**
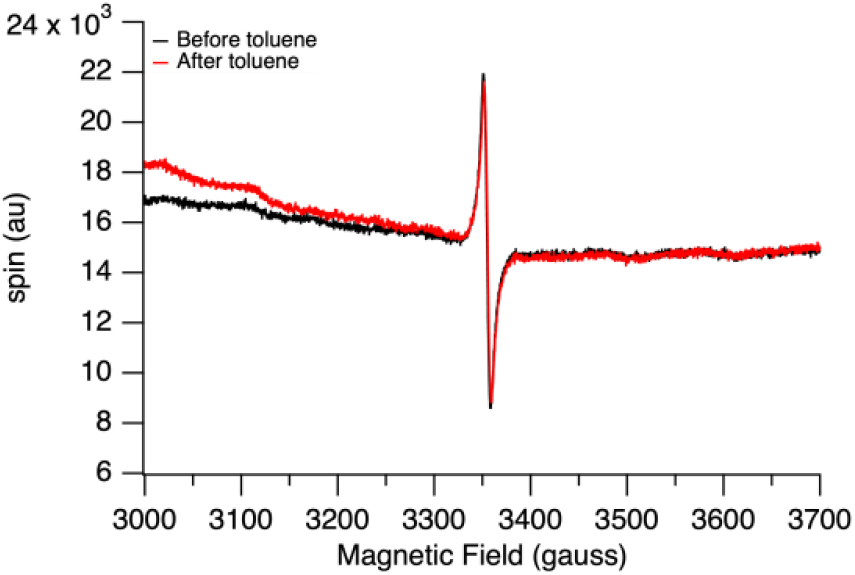
ESR spectra of live flies before and after exposure to toluene alone.

**Figure SM3:**
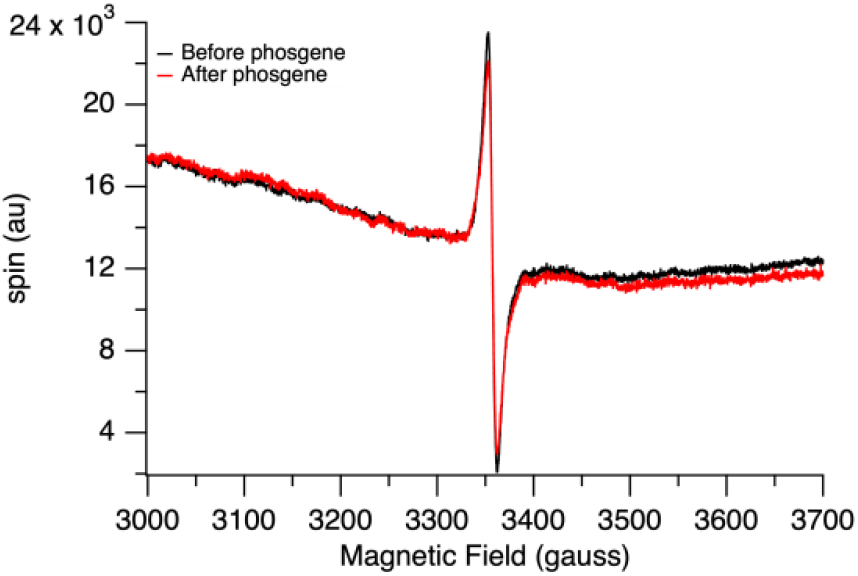
ESR spectra of dead flies before and after exposure to phosgene

**Figure SM5:**
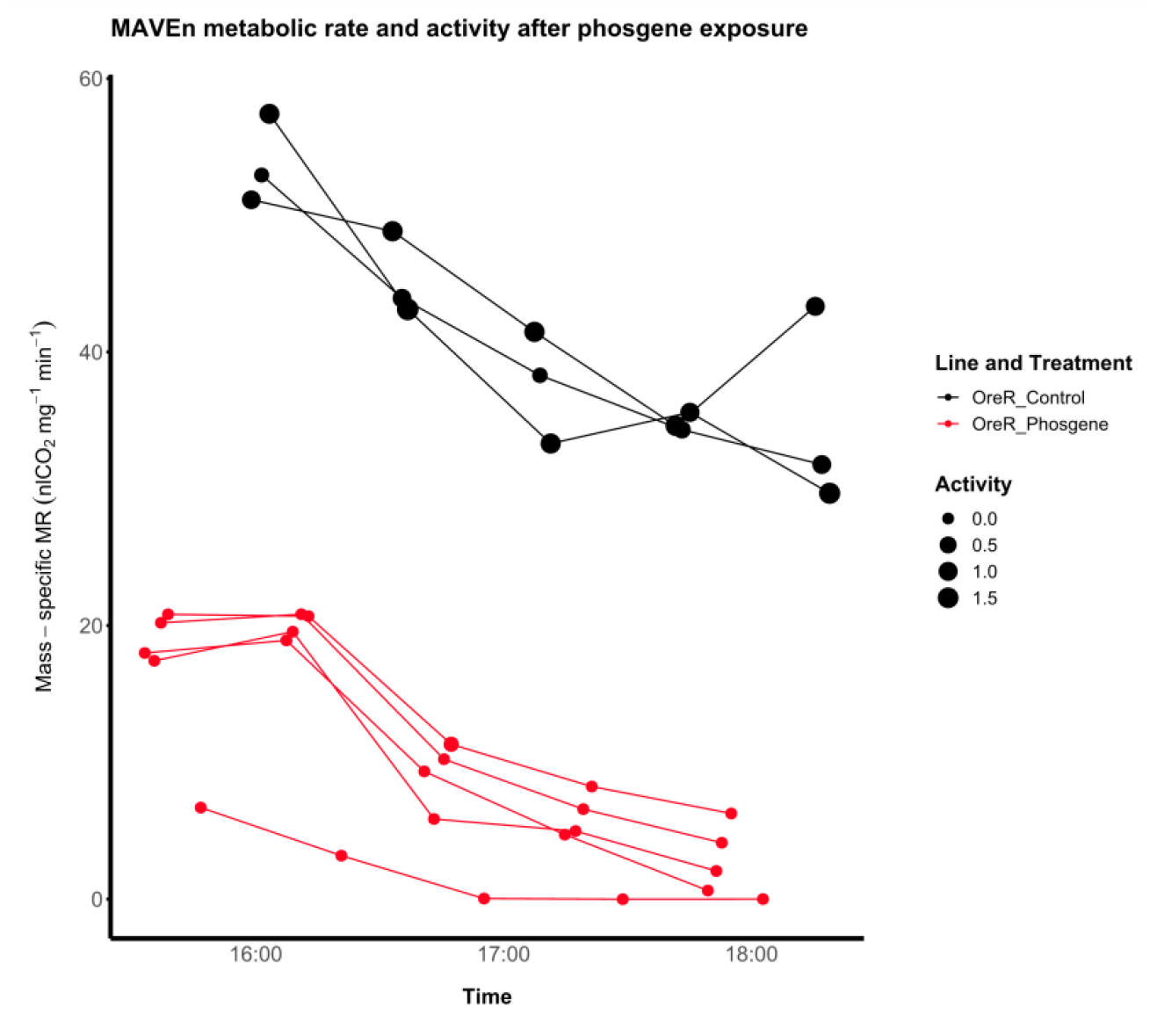
Multiple sequence alignment of MnSOD.

**Figure SM4:**
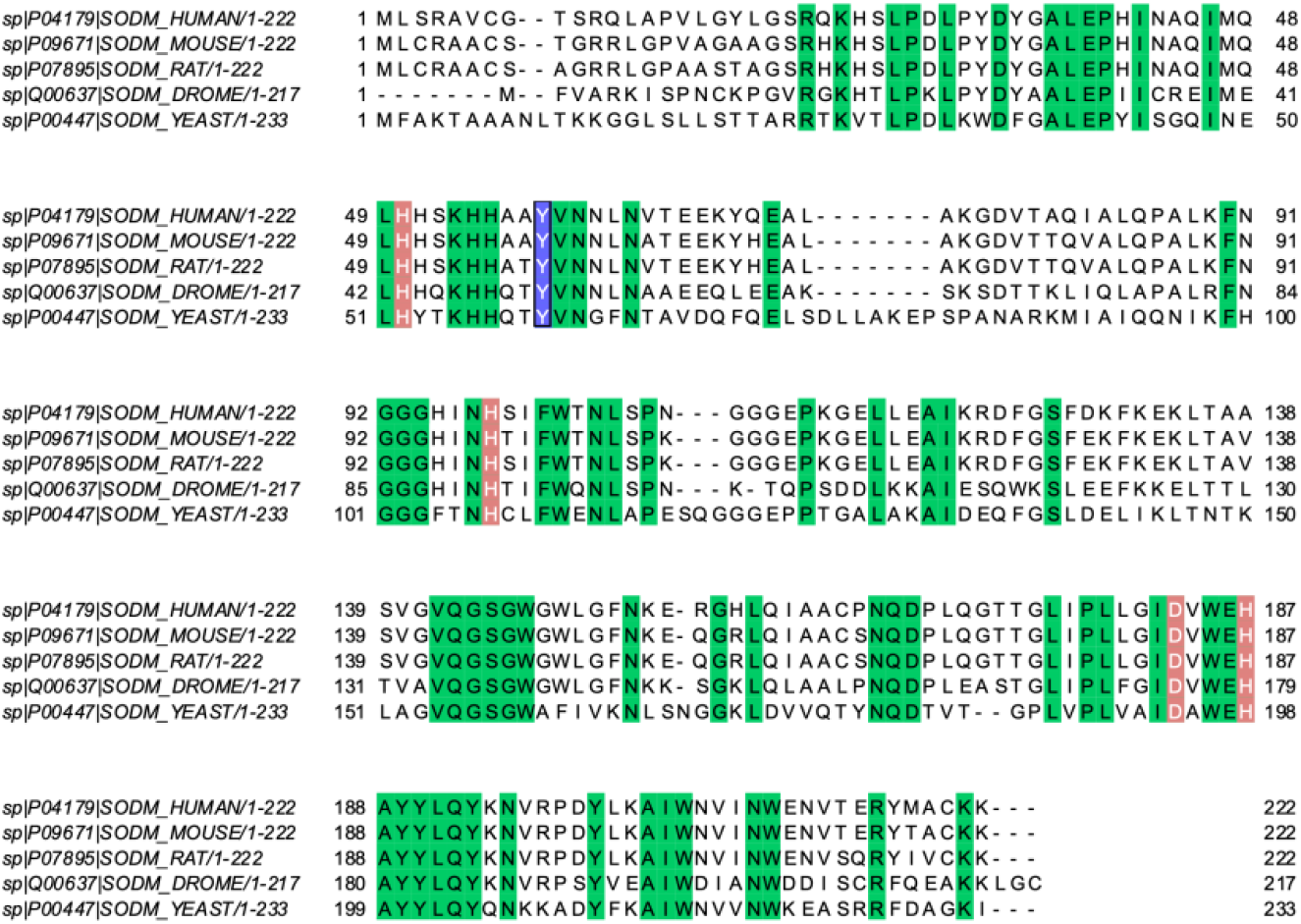
The mass-specific metabolic rate of flies was measured by Sable Instruments’ MAVEn on both untreated and treated flies with phosgene. Flies are weighted and placed inside the experiment tubes in groups of 5, and carbon dioxide consumption is measured through time.

## References

1. Turin, L., Skoulakis, E. M. C. & Horsfield, A. P. Electron spin changes during general anesthesia in Drosophila. Proc. Natl. Acad. Sci. U. S. A. 111, E3524–33 (2014).

2. Turin, L. & Skoulakis, E. M. C. Electron Spin Resonance (EPR) in Drosophila and General Anesthesia. Methods Enzymol. 603, 115–128 (2018).

3. Haffner, P. H., Goodsaid-Zalduondo, F. & Coleman, J. E. Electron spin resonance of manganese(II)-substituted zinc(II) metalloenzymes. J. Biol. Chem. 249, 6693–6695 (1974).

4. Babad, H. & Zeiler, A. G. Chemistry of phosgene. Chem. Rev. 73, 75–91 (1973).

5. Public Health Service, Agency for Toxic Substances and Disease Registry. Toxicological Profile for Chloroform. (1993).

6. Lawrence, G. D. & Sawyer, D. T. The chemistry of biological manganese. Coord. Chem. Rev. 27, 173–193 (1978).

7. Fridovich, I. Superoxide anion radical (O2-.), superoxide dismutases, and related matters. J. Biol. Chem. 272, 18515–18517 (1997).

8. Halliwell, B. & Gutteridge, J. M. C. Antioxidant defences synthesized in vivo. in Free Radicals in Biology and Medicine 77–152 (Oxford University Press, 2015).

9. Kallet, R. H. & Matthay, M. A. Hyperoxic acute lung injury. Respir. Care 58, 123–141 (2013).

10. Asikainen, T. M. et al. Increased sensitivity of homozygous Sod2 mutant mice to oxygen toxicity. Free Radic. Biol. Med. 32, 175–186 (2002).

11. Miriyala, S. et al. Manganese superoxide dismutase, MnSOD and its mimics. Biochim. Biophys. Acta 1822, 794–814 (2012).

12. Ho, Y. S., Vincent, R., Dey, M. S., Slot, J. W. & Crapo, J. D. Transgenic models for the study of lung antioxidant defense: enhanced manganese-containing superoxide dismutase activity gives partial protection to B6C3 hybrid mice exposed to hyperoxia. Am. J. Respir. Cell Mol. Biol. 18, 538–547 (1998).

13. Transcriptional Profiling of MnSOD-Mediated Lifespan Extension in Drosophilareveals a Species-General Network of Aging and Metabolic Genes.

14. Lebovitz, R. M. et al. Neurodegeneration, myocardial injury, and perinatal death in mitochondrial superoxide dismutase-deficient mice. Proc. Natl. Acad. Sci. U. S. A. 93, 9782–9787 (1996).

15. Alteration of Drosophila Life Span Using Conditional, Tissue-Specific Expression of Transgenes Triggered by Doxycycline or RU486/Mifepristone.

16. Pursuing the Elixir of Life: In Vivo Antioxidative Effects of Manganosalen Complexes.

17. Davy, J. VI. On a gaseous compound of carbonic oxide and chlorine. Philos. Trans. R. Soc. Lond. 102, 144–151 (1812).

18. Winternitz, M. C. & Yale University Anthony N Brady Memo. Collected Studies on the Pathology of War Gas Poisoning from the Department of Pathology and Bacteriology, Medical Science Section, Chemical Warfare Service, under the Direction of M.c. Winternitz. (Legare Street Press, 2021).

19. Nash, T. & Pattle, R. E. The absorption of phosgene by aqueous solutions and its relation to toxicity. Ann. Occup. Hyg. 14, 227–233 (1971).

20. Diller, W. F. Pathogenesis of phosgene poisoning. Toxicol. Ind. Health 1, 7–15 (1985).

21. Currie, W. D., Hatch, G. E. & Frosolono, M. F. Pulmonary alterations in rats due to acute phosgene inhalation. Fundam. Appl. Toxicol. 8, 107–114 (1987).

22. Pauluhn, J. Phosgene inhalation toxicity: Update on mechanisms and mechanism-based treatment strategies. Toxicology 450, 152682 (2021).

23. Hobson, S. T., Richieri, R. A. & Parseghian, M. H. Phosgene: toxicology, animal models, and medical countermeasures. Toxicol. Mech. Methods 31, 293–307 (2021).

24. Kent R. Olson Craig Smollin Ilene B. Anderson Neal L. Benowitz Paul D. Blanc Susan Y. Kim-Katz Justin C. Lewis Alan H.B. Wu. Poisoning & Drug Overdose. (2012).

25. Miller, F. J., Schlosser, P. M. & Janszen, D. B. Haber’s rule: a special case in a family of curves relating concentration and duration of exposure to a fixed level of response for a given endpoint. Toxicology 149, 21–34 (2000).

26. Toxicological Review of Phosgene. (2005).

27. Perry, J. J. P. et al. Contribution of human manganese superoxide dismutase tyrosine 34 to structure and catalysis. Biochemistry 48, 3417–3424 (2009).

28. Melov, S. et al. Mitochondrial disease in superoxide dismutase 2 mutant mice. Proc. Natl. Acad. Sci. U. S. A. 96, 846–851 (1999).

29. Campbell, J. B., Werkhoven, S. & Harrison, J. F. Metabolomics of anoxia tolerance in Drosophila melanogaster: evidence against substrate limitation and for roles of protective metabolites and paralytic hypometabolism. Am. J. Physiol. Regul. Integr. Comp. Physiol. 317, R442–R450 (2019).

30. Camus, M. F., Rodriguez, E., Kotiadis, V., Carter, H. & Lane, N. Redox stress shortens lifespan through suppression of respiratory complex I in flies with mitonuclear incompatibilities. Exp. Gerontol. 175, 112158 (2023).

31. Rodríguez, E., Bettinazzi, S., Inwongwan, S., Camus, M. F. & Lane, N. Harmonizing protocols to measure Drosophila respiratory function in mitochondrial preparations. (2023) doi:10.26124/BEC:2023-0003.

32. Correa, C. C., Aw, W. C., Melvin, R. G., Pichaud, N. & Ballard, J. W. O. Mitochondrial DNA variants influence mitochondrial bioenergetics in Drosophila melanogaster. Mitochondrion 12, 459–464 (2012).

33. Trapp, C., Waters, B., Lebendiger, G. & Perkins, M. Electron spin resonances of a living system (Drosophila) on normal and carcinogenic diets. Biochem. Biophys. Res. Commun. 112, 602–605 (1983).

34. Kar, A. K., Acharya, A., Pradhan, G. C. & Dash, A. C. Glyoxylate as a reducing agent for manganese(III) in salen scaffold: A kinetics and mechanistic study. J. Chem. Sci. (Bangalore) 126, 547–559 (2014).

35. FlyBase. FlyBase gene report: Dmel\Sod2. http://flybase.org/reports/FBgn0010213.htm.

36. Azadmanesh, J. & Borgstahl, G. A review of the catalytic mechanism of human manganese superoxide dismutase. Antioxidants (Basel) 7, 25 (2018).

37. Matzner, M., Kurkjy, R. P. & Cotter, R. J. The chemistry of chloroformates. Chem. Rev. 64, 645–687 (1964).

38. Wang, Y., Branicky, R., Noë, A. & Hekimi, S. Superoxide dismutases: Dual roles in controlling ROS damage and regulating ROS signaling. J. Cell Biol. 217, 1915–1928 (2018).

39. Holmes, W. W. et al. Conceptual approaches for treatment of phosgene inhalation-induced lung injury. Toxicol. Lett. 244, 8–20 (2016).

40. Borak, J. & Diller, W. F. Phosgene exposure: mechanisms of injury and treatment strategies. J. Occup. Environ. Med. 43, 110–119 (2001).

41. Bruner, H. D., Boche, R. D., Chapple, C. C., Gibbon, M. H. & McCarthy, M. D. Studies on experimental phosgene poisoning. Iii. Oxygen therapy in phosgene-poisoned dogs and rats. J. Clin. Invest. 26, 936–944 (1947).

42. Hough, L. & Phadnis, S. P. Enhancement in the sweetness of sucrose. Nature 263, 800 (1976).

43. Asgari, A., Parak, M., Nourian, Y. H. & Ghanei, M. Phosgene toxicity clinical manifestations and treatment: A systematic review. Cell J. 26, 91–97 (2024).

44. Pawlowski, R. & Frosolono, M. F. Effect of phosgene on rat lungs after single High-Level exposure. Arch. Environ. Health 32, 278–283 (1977).

45. Sciuto, A. Inhalation toxicology of an irritant gas© 2006 by Taylor & Francis group, LLC historical perspectives, current research, and case studies of phosgene exposure. in Inhalation Toxicology, Second Edition 457–483 (CRC Press, 2005).

46. Wong, B., Perkins, M. W. & Sciuto, A. M. Phosgene. in Hamilton & Hardy’s Industrial Toxicology 357–362 (John Wiley & Sons, Inc., Hoboken, New Jersey, 2015).

47. Lu, Q. et al. Mechanism of phosgene-induced acute lung injury and treatment strategy. Int. J. Mol. Sci. 22, 10933 (2021).

48. Jaskot, R. H., Grose, E. C., Richards, J. H. & Doerfler, D. L. Effects of inhaled phosgene on rat lung antioxidant systems. Toxicol. Sci. 17, 666–674 (1991).

49. Clark, J. M. The toxicity of oxygen. Am. Rev. Respir. Dis. 110, 40–50 (1974).

50. Thomson, L. & Paton, J. Oxygen toxicity. Paediatr. Respir. Rev. 15, 120–123 (2014).

51. McCord, J. M. & Fridovich, I. The biology and pathology of oxygen radicals. Ann. Intern. Med. 89, 122–127 (1978).

52. Holley, A. K., Bakthavatchalu, V., Velez-Roman, J. M. & St Clair, D. K. Manganese superoxide dismutase: guardian of the powerhouse. Int. J. Mol. Sci. 12, 7114–7162 (2011).

53. Stallings, W. C., Pattridge, K. A., Strong, R. K. & Ludwig, M. L. The structure of manganese superoxide dismutase from Thermus thermophilus HB8 at 2.4-A resolution. J. Biol. Chem. 260, 16424–16432 (1985).

